# Vasoactive Endothelial Growth Factor and Heat Shock Protein Gene Expression Response in Kawasaki Disease

**DOI:** 10.1101/2022.09.26.508411

**Authors:** Asrar Rashid, Hoda Alkhzaimi, Govind Benakatti, Zainab A. Malik, Varun Sharma, Anuka Sharma, Rayaz Malik, Nasir Quraishi, Guftar Shaikh, Ahmed Al-Dubai, Amir Hussain

## Abstract

Kawasaki Disease (KD) is a childhood vasculitis primarily affecting medium-sized arteries, which can lead to severe complications, particularly with respect to coronary artery disease (CAD). The impact of thermal stress on KD coronary artery pathogenesis, in association with prolonged fever and inflammation, remains unclear. In this study, we hypothesized that altered gene expression (GE) of angiogenesis-inducing Heat Shock Proteins (HSPs) is associated with KD-CAD through pro-inflammation. Transcriptomic analysis was performed using the three largest KD peripheral blood studies in the clinical literature (KD1-KD3), and one study direct from coronary artery tissue (KD4). The analysis revealed a significant increase in TNF and NFKB1 GE, indicating the presence of inflammation based on gene expression profiles. Gene set enrichment analysis (GSEA) of KD1-KD3 datasets identified inflammatory pathways, including TNFA signaling via NFKB, IL6 JAK STAT 3 Signalling, and p53 (Heat Shock Protein 90). The study also focused on specific HSPs known to be associated with angiogenesis, namely HSPB1, HSPA1A, and HSP90AB1. The temporal transcript model (TTM) consistently showed up-regulation of pro-inflammatory genes VEGF-A, TNF, and NFKB1, as well as up-regulation of HSPA1A. GSEA revealed gene ontology pathways associated with VEGF production. These findings suggest that the binding of VEGF-A or VEGF-B to their receptors could potentially impact the coronary artery in KD. Additionally, the up-regulation of the gene HSPAB1 in KD has not been described previously. In contrast, KD4 showed no differential GE for the studied genes potentially related to end-stage KD. This study provides valuable insights into VEGF and HSPs in KD-associated inflammation. Future research should focus on developing a VEGF-HSP CAD model to explore implications for KD biomarking as well as developing precision management strategies.

## Introduction

Kawasaki Disease (KD) is the leading cause of acute childhood vasculitis affecting medium-sized arteries, with a higher incidence in deprived socioeconomic groups^1^. KD predominantly affects coronary arteries with a risk of coronary artery disease (CAD) if left untreated. Early recognition is crucial, but diagnosis remains challenging due to the overlap of symptoms with common childhood febrile illnesses. KD diagnostic criteria include a fever lasting for five or more days as a key feature and associated with a period of intense inflammation, consistent with increased CRP and Pro-calcitonin^2^. A prospective study of 60 children revealed increased pro-inflammatory cytokines IL-6 and IL-8 during the first week of KD, with higher levels differentiating children who developed coronary artery aneurysms^3^. Then, in the second week of KD, the formation of coronary aneurysms was associated with elevated TNF levels. Pro-inflammatory cytokines like IL-6, IL-8, and TNF enhance the immune response by triggering further inflammation.

How a persistent fever and inflammation in relation to KD translates to coronary disease pathogenesis remains largely unknown. However, more is known about thermal stress mechanisms associated with the protective action of Heat shock proteins (HSPs) outside the context of KD. HSPs protect the host from thermal stress during heightened inflammation being ubiquitous chaperone proteins that provide cellular quality control and assist the correction of misfolded nascent proteins^4^. Several experimental studies confirm the association between HSP and angiogenesis ^5–7^. HSPs with functionality related to angiogenesis are shown (Table 1). Additionally, the family of proteins known as Vascular Endothelial Growth Factors could also have a role in KD CAD in association with HSPs. Elevated VEGF levels have been found in acute KD patients with and without coronary pathology^8^. The best studied in KD is VEGF-A, which is pro-inflammatory and is associated with worse outcomes in ST-segment myocardial infarction ^9^. Another sub-type, VEGF-B, promotes cell survival and migration of endothelial cells, though its exact role in KD has not been substantiated.

**TABLE 1:** Heat Shock Proteins with a Role in Angiogenesis

|  | Family membership | Full Name | Associated Function | Ref |
| --- | --- | --- | --- | --- |
| **HSPB1 (HSP27)** | HSP20 | Heat shock protein family B (small) member 1 | Cell growth and differentiation owing to dual antioxidant and protein chaperone properties |  |
|  |  |  | HSPB1 protects cells from various stressors, including heat and ischemia | 30 |
|  |  |  | The role of HSPB1 may not be specific, as HSPB1 is increased in non-infective stressful states, such as due to post-trauma-induced inflammation | 31 |
|  |  |  | HSPB1-VEGF interactions may be important in the balance between physiological and pathological angiogenesis | 17 |
| **HSPA1A** | HSP70 | HSP70 Family 1A | Has both angiogenic and chaperone properties by inducing endothelial cell migration and micro-vessel formation | 32 |
| **HSP90AB1** | HSP90 | Heat shock protein 90 alpha family class B member 1 | HSP90AB1 is a heat shock protein that has also been implicated in regulating several signaling pathways involved in angiogenesis, such as the VEGF pathway.<br>Studies suggest HSP90AB1 acts as a "chaperone" for VEGF receptors and other signaling proteins.<br>HSP90AB1 stabilizes VEGF receptors and enhances their signaling capacity, leading to the stimulation of angiogenesis. Inhibitors of HSP90AB1 have been shown to reduce angiogenesis in preclinical studies, suggesting that HSP90AB1 plays an important role in this process. | 33 |
| **HSPBAP1** | Binds to HSP27 | HSPB1 associated protein 1 | HSPBAP1 has been associated with two rare angiogenesis-related conditions (Orpha ID 217071 and 151)<br>The HSPBAP1 gene encodes a protein binding to the small heat shock protein Hsp27. Hsp27 is involved with cell growth and differentiation.<br>Role in conditions such as KD is unknown | 34 |

In previous work in KD, we have highlighted differential gene expression (GE) for genes associated with pro-inflammatory proteins (TNF and NFKB1), pro-angiogenic HSPs, and VEGF (VEGF-A and VEGF-B) ^10^. However, the function of HSPs in KD, and particularly the inter-relationship between HSPs and VEGF in KD pathogenesis during the persistent febrile response, has received little attention in the clinical literature. The primary aim of this study was to assess KD-associated gene pathways and differential gene expression (GE) for genes associated with pro-inflammatory proteins (TNF and NFKB1), pro-angiogenic HSPs, and VEGF (VEGF-A and VEGF-B).

## Material and Methods

### Using a Relevant Framework for the Study

To research the association of VEGF, HSP, and angiogenesis, a VEGF-HSP framework was adopted, entailing a Kyoto Encyclopedia of Genes and Genomes (KEGG) P13K-Akt signaling pathway (identifier 04370 8/28/17) (Figure 1). This KEGG pathway outlines the link between VEGF and endothelial migration and proliferation. Further, the map shows that HSPB1 (HSP27) and VEGF-A share a common pathway responsible for inflammation, endothelial migration, and endothelial proliferation. Moreover, this diagram illustrates that the VEGF receptor 2 (VEGFR2) pathway is mediated by VEGF-A acting on the endothelium, suggesting the potential importance of HSPB1 in angiogenesis. VEGFR-2 is an essential transducer in angiogenesis in physiological and pathological conditions ^11^.

**Figure 1.**
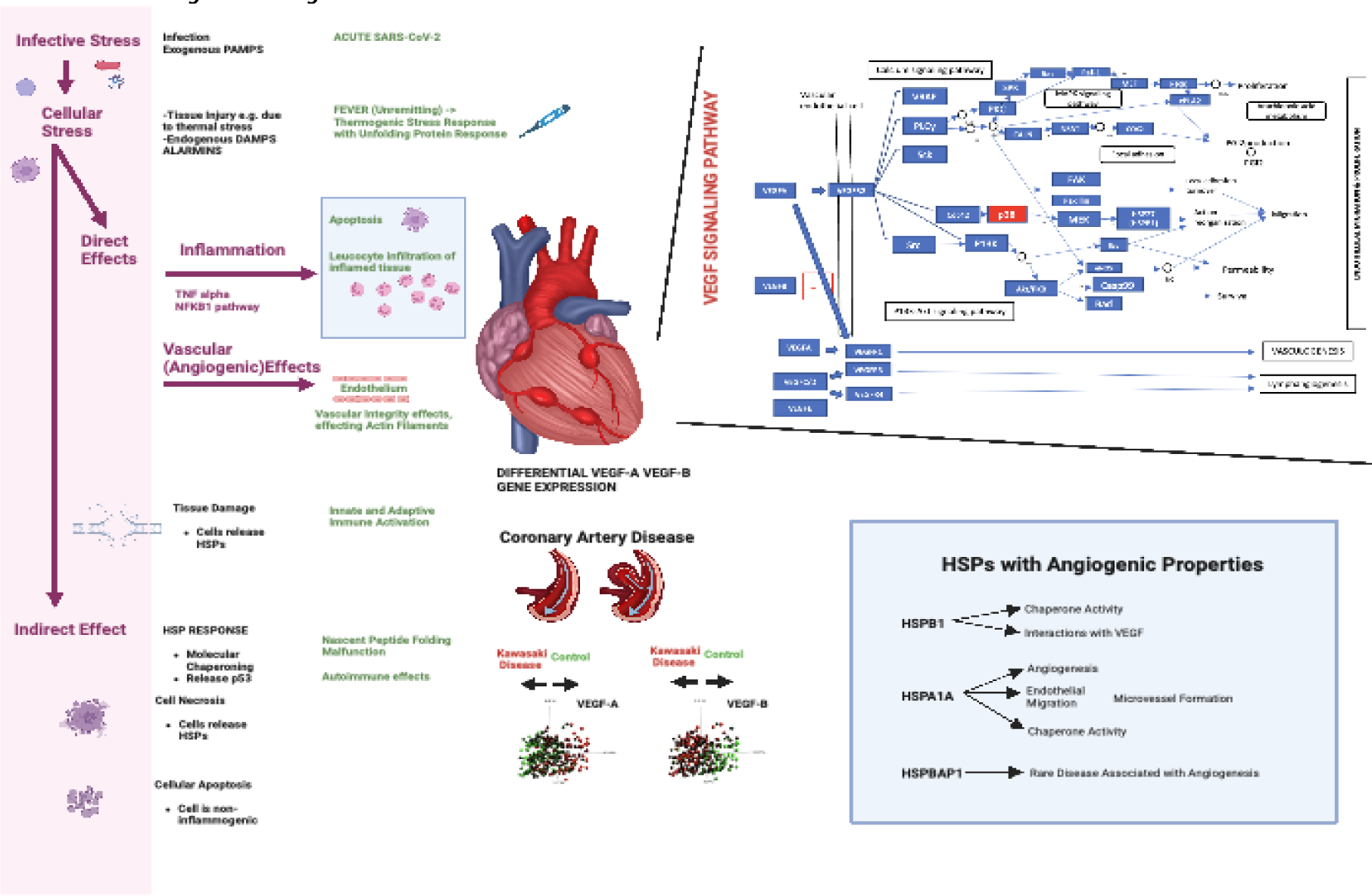
Mechanisms of CAD with the HSP-VEGF mechanisms lead to endotheliitis and coronary aneurysm formation. Figure 1. The etiology of KD remains unknown, but there is a possibility this may be associated with an infected agent (upper left of diagram), as exemplified with the SARS-CoV2 pandemic and the development of a KD-like illness known as Multi-inflammatory Disease in Childhood (MIS-C). The left side of the diagram portrays the relationship of a primary infectious stressor leading to thermal stress, which in turn initiates direct and indirect immune activation. The capability to distinguish infected host cells is mediated through pattern recognition receptors (PRRs) on lymphocytes and antigen-presenting cells (left figure)^28^. Pathogen-associated molecular patterns (PAMPs) may be present in pathogenic cells and their derivatives. The concept of danger signals has evolved to encompass both endogenous and exogenous triggers, where host products can stimulate a danger trigger, playing the role of an alarmin. How an acute infective agent might lead to KD is not known. Also, how the KD evolves from the acute stage to convalescence from the molecular point of view is also not known. KD culminates in a medium vessel arteritis with a predisposition in children for the coronary architecture resulting in aneurysm formation and mortality risk. The figure in the upper right, adapted from a KEGG Vascular Endothelial Growth Factor (VEGF) signaling pathway (KEGG pathway map identifier 04370 8/28/17), indicates mechanisms integral to the pathogenesis of KD as per KEGG nomenclature. It illustrates VEGF-A binding to VEGFR2 on the vascular endothelial cell surface. VEGF-A’s action on the endothelium could play a role in the formation of coronary artery lesions and aneurysms in KD. As represented in the diagram, VEGF-A’s impact via the PI13k-Akt signaling pathway triggering various outcomes, including endothelial migration and angiogenesis^29^. The role of other VEGF subtypes in KD has not been investigated in detail in the scientific literature. Acute KD changes includes extracellular leakage from cell injury and necrosis, potentially involving Heat Shock Protein (HSP)-associated processes. HSPs play a key role in mechanisms related to thermal stress and in the quality control of newly folded proteins. Protein-folding malfunctions could potentially trigger various immune activation phenomena, including autoimmunity. The inflammatory cascade culminating in endothelial migration and dysangiogenesis, can involve proteins such as HSPB1 and an HSP may be related to VEGF-A stimulation. The influence of HSP genes (HSPB1, HSPBAP1 and HSPA1A) on angiogenesis is displayed in the lower right quadrant of the diagram. This pathway might provide insights into why anti-inflammatory treatments don’t impact endothelial effects and the associated inflammation resulting in coronary arteritis. An unremitting fever is characteristic of KD, which is why it is included in the diagnostic criteria. Standard anti pyretics and anti-inflammatories, such as acetaminophen and non-steroidal drugs are utilized to manage fever in children with KD. Why these are inadequate in limiting KD associated inflammation is explainable by reference to the aforementioned KEGG pathway. These drugs operate via separate PGI2 pathways affecting COX-2-based mechanisms. Here host products may stimulate a danger trigger, a role known as being an alarmin. The final figure was finalised and created with BioRender.com

### Dataset Inclusion and Search Strategy

Datasets were obtained from the National Centre for Biotechnology Information (NCBI; https://www.ncbi.nlm.nih.gov) Gene Expression Omnibus (GEO, http://www.ncbi.nlm.nih.gov/geo/) (Table 2). To ensure a robust foundation for drawing inferences about gene expression patterns, studies with a large sample size were sought. An arbitrary threshold of greater than 100 samples per study was chosen, which excluded studies the other studies (which ranged from n=9 to 41). Three KD datasets (KD1:GSE63881, KD2:73464, and KD3:GSE68004) were thus selected based on a search strategy (Figure 6). These datasets coincided with the datasets from our earlier work on KD, showing the datasets to elicit information important in understanding KD pathogenesis^10^. Additionally, a unique dataset (GSE64486), believed to be the only one derived from the coronary arteries of children with KD, was also included in the analysis.

**TABLE 2:** Kawasaki Disease Datasets Selected for Analysis

| Accession number* | Platform | Named in the body of text | Species | Disease | STUDY DESIGN | Subject Number | Ref*** |
| --- | --- | --- | --- | --- | --- | --- | --- |
| GSE63881 | Illumina HumanHT-12 V4.0 beadchip | Microarray Study | Human | KD1 | Acute KD (n=171) Versus Convalescent (n=170) samples | 146 | {Hoang, 2014 #2320} |
| **GSE73461 | Illumina HumanHT-12 V3.0 beadchip | Microarray Study | Human | KD2 | KD (n=78) versus Control (n=55) | 133 | {Wright, 2018 #2323} |
| **GSE7363 | Illumina HumanHT-12 V3.0 beadchip | Microarray Study | Human | KD2 | Acute KD (n=146) Versus Convalescent Samples (n=87) | 233 | {Wright, 2018 #2323} |
| GSE68004 | Illumina HumanHT-12 V4.0 beadchip | Microarray Study | Human | KD3 | KD (n=89) versus Controls 37 | 126 | {Jaggi, 2018 #2321} |
| GSE64486 | Illumina HiSeq 2000 | RNA-Seq | Human | KD4 | Coronary artery tissue from KD (n=8) and Childhood controls (n=7) | 15 | {Rowley, 2015 #2394} |

\* NCBI GEO Accession Number or NCBI Bioproject ID is given
\*\* GSE73464 is the super series containing sub-series studies GSE73461 and GSE73463, 73462 was not analysed as it contained only one case of KD.
\*\*\*Referebnce

### Microarray Dataset Pre-processing

All microarray datasets were pre-processed by log2 quantile normalization using R-script.

### RNA-Seq Datasets

The paired-end RNAseq FASTQ files for each sample within the dataset GSE173317 were obtained using the tool parallel-fastq-dump (version 0.67, fastq-dump version 2.8.0) and the reads were mapped to the human reference genome (hg38 obtained from UCSC, URL: https://hgdownload.soe.ucsc.edu/goldenPath/hg38/bigZips/hg38.fa.gz) using the BWA-MEM algorithm (bwa version 0.7.17-r1188). Finally, the resulting SAM files, containing the mapped reads, were sorted by coordinate using Picard - SortSam (version 2.25.5) to generate mapped BAM files from each sample. A Gene Transfer Format (GTF) file was generated from the UCSC.edu table browser using the settings clade ‘Mammal,’ genome ‘Human’ assembly Dec.2013 (GRCh38/hg38), group ‘Genes and Gene Predictions,’ track ’Gencode V36’, table ‘knownGene”. The BAM files created were analyzed in the TMM setting of QOE.

### Study Design

The selected studies were either case-controlled, comparing KD patients with controls, or examined the acute phase of KD against convalescence (Table 2). Genes were identified through bioinformatics analysis, and different subgroups within each KD cohort were utilized to understand significant gene sets. To investigate inflammation patterns, the goal was to define alterations in gene expression patterns associated with specific genes in whom corresponding proteins are known to be pro-inflammatory, such as TNF, NFKB1, and VEGF-A. VEGF-B was also chosen as a comparison against VEGF-A, as both proteins compete for the same receptor, VEGF-R1. HSPs were chosen based on their known associations with angiogenesis.

## Bioinformatics Software

### Statistical and Gene Ontology analyses

Differential expression of the genes (DEGs) was undertaken using Qlucore Omics Explorer (QOE) version 3.1 software (Qlucore AB, Lund, Sweden). Principal Component Analysis (PCA) plots were generated from QOE. This allowed two-group and multi-group comparisons and unsupervised hierarchical clustering depicted by heat plots. mRNA array data was analyzed with Qlucore Omics Explorer version 3.7 for DEGs. To understand the genes of interest, Gene Symbols were generated for the Genes from the data to compensate for the multiple probes using an averaging method and applied according to the analysis. Hierarchical clustering was based on Euclidean distance and average linkage clustering, and all genes were centered to a mean equal to zero and scaled to variance similar to one. Value q, also known as the False discovery rate (FDR), was used to support the raw p-value. A p-value less than 0.05 and FDR (q) value below 0.05 was considered statistically significant for the statistical aspect of this paper. QOE was also used to generate a PCA heat plot of genes of interest. Bulk tissue expression of VEGFA and VEGFB was performed using the GTEx biobank portal, and Transcripts Per Million (TPM) for RNA-seq values were recorded on the tissues associated with cardiac associated tissues (https://gtexportal.org/) using GTEx Analysis Release V8 (dbGaP Accession phs000424.v8.p2). Moreover, a gene function network interaction was performed between VEGF (VEGF-A, VEGF-B, VEGF-C and VEGF-D) and HSP genes (HSPBAP1 and HSPB1) using the GeneMANIA platform^12^.

### Gene Set Enrichment Analysis

Gene Set Enrichment Analysis (GSEA) was undertaken using the QOE platform, comparing three KD studies. For KD1 acute versus convalescent for KD2 and KD3 versus controls. To undertake a GSEA comparison, gene sets were downloaded from the MSigDB database generating GMT files. GMT files were labeled according to the specific search terms in the database, such as ‘apoptosis,’ ‘CD40,’ ‘VEGF,’ and so on. The Hallmark GMT file was generated from version 7.2 of the Hallmark geneset downloaded from the MSigDB database.

### Actin Analysis (GSEA)

KD1-KD3 were analyzed using actin-related gene sets to gain insights into the function of actin in KD pathogenesis.

### Differential Gene Expression Analysis

For microarray analysis, HSPs chosen for gene expression analysis based on a relationship to angiogenesis included Heat Shock Protein Beta 1 (HSPB1), HSPA1A, HSP27, and HSPBAP1. Genes associated with inflammation in KD were selected, including TNF and NFKB1, and in relation to VEGF, gene expression for VEGF-A and VEGF-B was included.

## Results

### A. TNF and NFKB1 GE (KD1-KD3)

TNF and NFKB1 GE was compared across KD studies (Fig.2 G-I #A-C).

**Figure 2.**
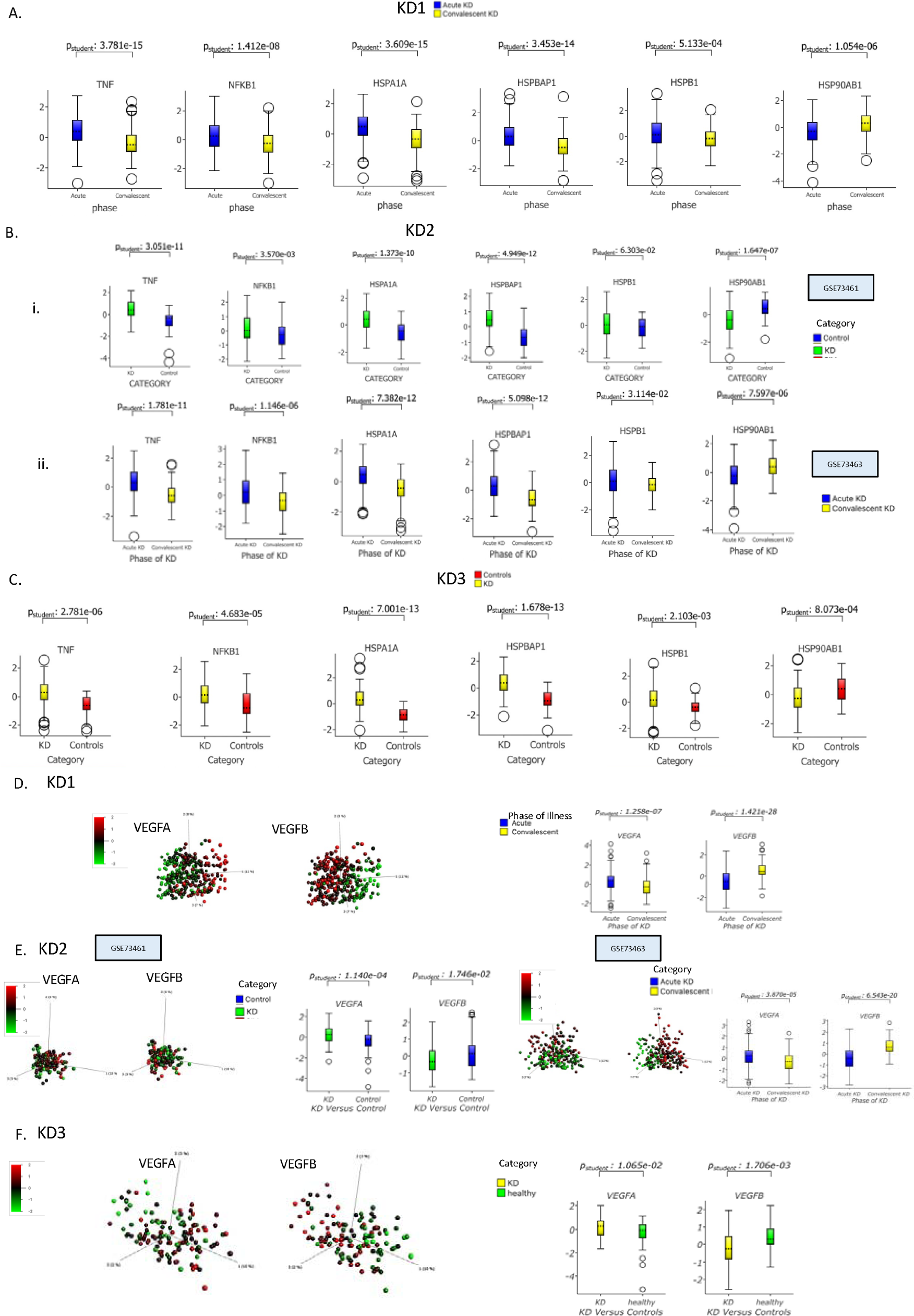
VEGFA, VEGFB, HSP’s, TNF, NFKB1 Gene-Expression in KD datasets. **Figure 2.** Kawasaki Dataset PCA heat map gene expression and box plots are shown, KD1 (Figures 2A & B), KD2 (Figures 2C & D), and KD3 (Figures 2E & F). For KD1 acute versus convalescent samples, KD2 and KD3 (KD versus control samples), there is a significant downregulation of VEGFB, HSPA1A, HSPBAP1, TNF, and NFKB1 gene expression. HSPB1 gene expression did not differ in KD2 but was increased in acute versus convalescent samples and KD3 compared to control samples. In KD1 (acute versus convalescent samples), KD2, and KD3 (KD versus control samples), there is significant up-regulation in VEGFA gene expression. In acute versus convalescent samples (KD1), VEGF A is up-regulated (p=1.258e-07), VEGF B is down-regulated (p=1.42e-28), HSP90AB1 is down-regulated (p=1.054e-06), and HSPBAP1 is up-regulated (p=3.453e-14). In KD versus control (KD2), VEGFA is up-regulated (p=1.140e-04), VEGFB down-regulated (p=1.746e-02), HSP90AB1 down-regulated (p=1.647e-07), HSPBAP1 up-regulated (p=4.949e12). In KD versus control (KD3), VEGFA is up-regulated (p=1.140e-04), VEGFB down-regulated (p=1.746e-02), HSP90AB1 down-regulated (p=1.647e-07), and HSPBAP1 up-regulated (p=4.949e-12).

**Figure 3.**
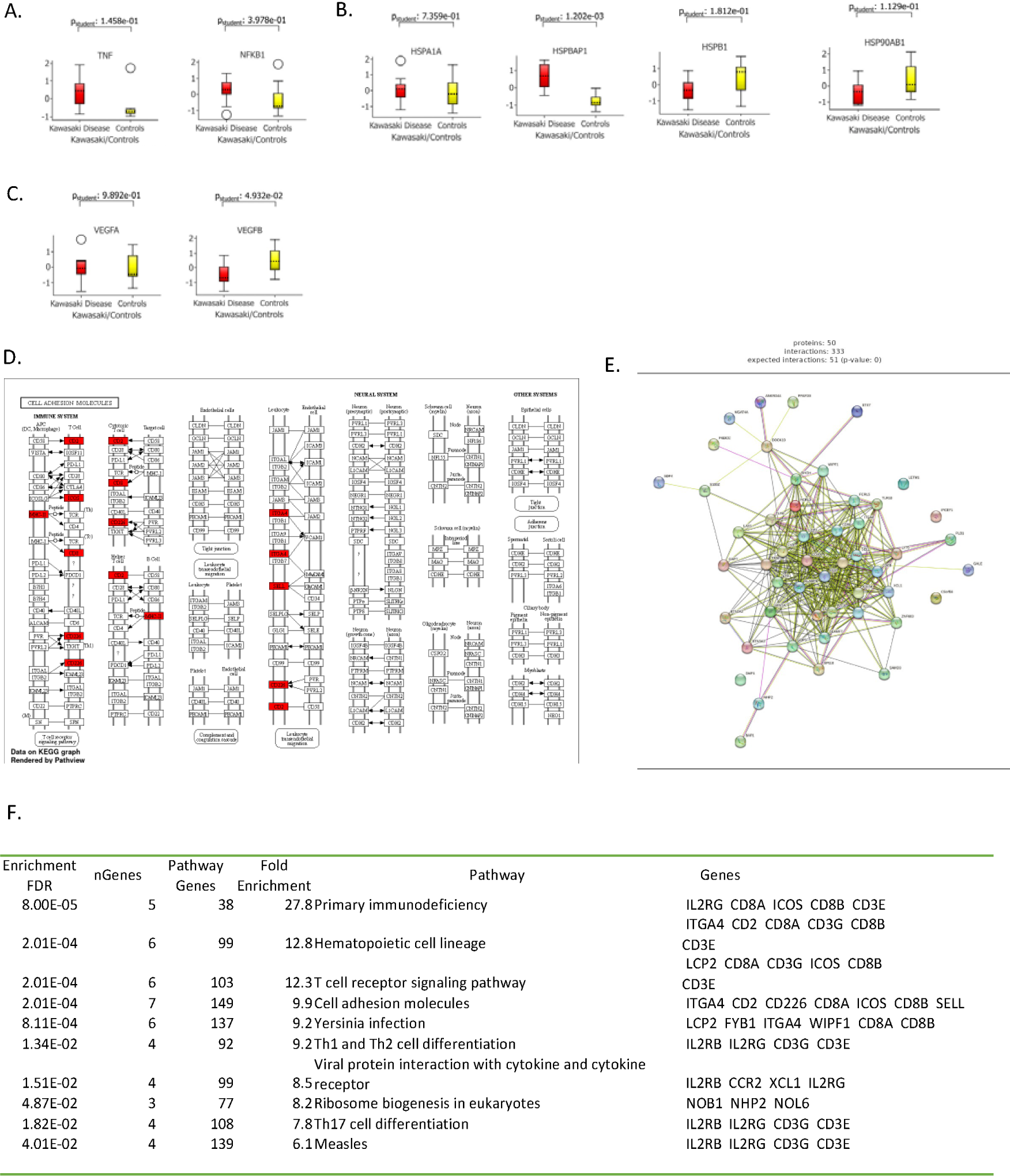
Coronary Artery Gene Expression in KD with Coronary Artery Disease Compared to Controls. **Figure 3.** Analysis of coronary artery gene expression based on the KD4 (GSE64486) dataset comparing KD patients and controls. TNF and NFKB1 gene expressions (GE) exhibited no significant differences between the two groups (Figure 3A). Similarly, no differential GE was observed for the studied heat shock proteins (HSPA1A, HSPBAP1, HSPB1, and HSP90AB1)(Figure 3B), or for VEGF-A to VEGF-B (Figure 3C). Excluding infected controls from the analysis did not enhance GE differentiation. Adjusting the PCA plot (p=0.001 and q=0.12) highlighted 192 genes that subsequently underwent enrichment analysis. This process led to the creation of the KEGG (cell adhesion) diagram via Pathview (Figure 3D), with enriched genes depicted in red. These 192 genes were processed through the Shinygo platform and submitted to the STRING-db website for additional enrichment analysis, resulting in a protein-protein interaction network. The data was mapped against human, archaeal, bacterial, and eukaryotic species in the STRING server database, successfully mapping 55.7% of the genes (Figure 3E). Ten of the most significantly enriched genes are displayed (Figure 3F).

TNF and NFKB1 GE was increased in Acute Versus convalescent samples [KD1: p=3.781e-15, p=1.412e-08 and KD2: p=1.781e-11, p=1.148e-06] and KD versus controls [KD2: p=3.051e-11, KD3: p=3.570e-03].

### B. HSP Response in Acute KD (KD1-KD3)

Gene expression for HSPs associated with angiogenesis (HSPA1A, HSPBAP1, HSB1, HSP90AB1) was also analyzed through box plot analysis (Fig 2 A-C). In all three KD datasets, HSPA1A and HSBAP1, HSPB1 genes were up-regulated in acute versus convalescent samples [KD1: p=3.609e-15, p=3.453e-14, p=5.133e-04 and for KD2: p=7.382e-12, p=5.098e-12, p=3.114e-02]. For KD versus controls genes HSPA1A, HSPBAP1 were up-regulated [KD2: p=1.373e-10, p=4.949e-12 and KD3: p=7.001e-13, p=1.678e-13]. However, for KD versus controls, HSPB1 GE in KD showed no difference [KD2: p=6.303e-02] and was also up-regulated [KD3: p=2.103e-03]. For HSP90AB1 GE showed downregulation in Acute Versus convalescent samples [KD1: p=1.054e-06, KD2: p=7.597e-06] and KD versus controls [KD2: p=1.647e-07, KD3: p=8.073e-04].

### C. VEGF GE (KD1-KD3)

VEGF-A and VEGF-B GE patterns were compared across the studies (KD1-D3) using heat-map and box plot analysis (Fig.2 A-E #D-F). VEGF-A GE was increased in Acute Versus convalescent samples [KD1: p=1.258e-07, KD2: p=3.870e-05] and KD versus controls [KD2: p=1.140e-04, KD3: p=1.065e-02] VEGF-B GE was decreased in Acute Versus convalescent samples [KD1: p=1.421e-28, KD2: p=6.543e-20], in KD versus controls this was decreased [KD2: p=1.746e-02, KD3: p=1.706e-03]

### D. Coronary Arteritis (KD4: GSE64486)

Coronary artery samples were analyzed against controls. In cases of KD with versus controls, GE for TNF and NFKB1 showed no difference (p=1.458e-01, p=3.978e-01). For the HSP’s: HSPA1a, HSPBAP1, HSPB1 and HSP90AB1, again there was no difference in GE (p=7.359e-01, p=1.202e-03, p=1.812e-01 and p=1.129e-01). For VEGF-A, and VEGF-B there was also no difference between KD and controls (p=9.892e-01, p=4.932e-02, p=1.585e-01, p=3.473e-01).

### E. GSEA (KD1-KD3)

Table 3 shows the results of GSEA performed on acute versus convalescent samples (KD1) and KD versus control samples (KD2 and KD3), using different reference sets to analyze the significance of various pathways. The analysis revealed several noteworthy themes.

**Supplement TABLE 3:**
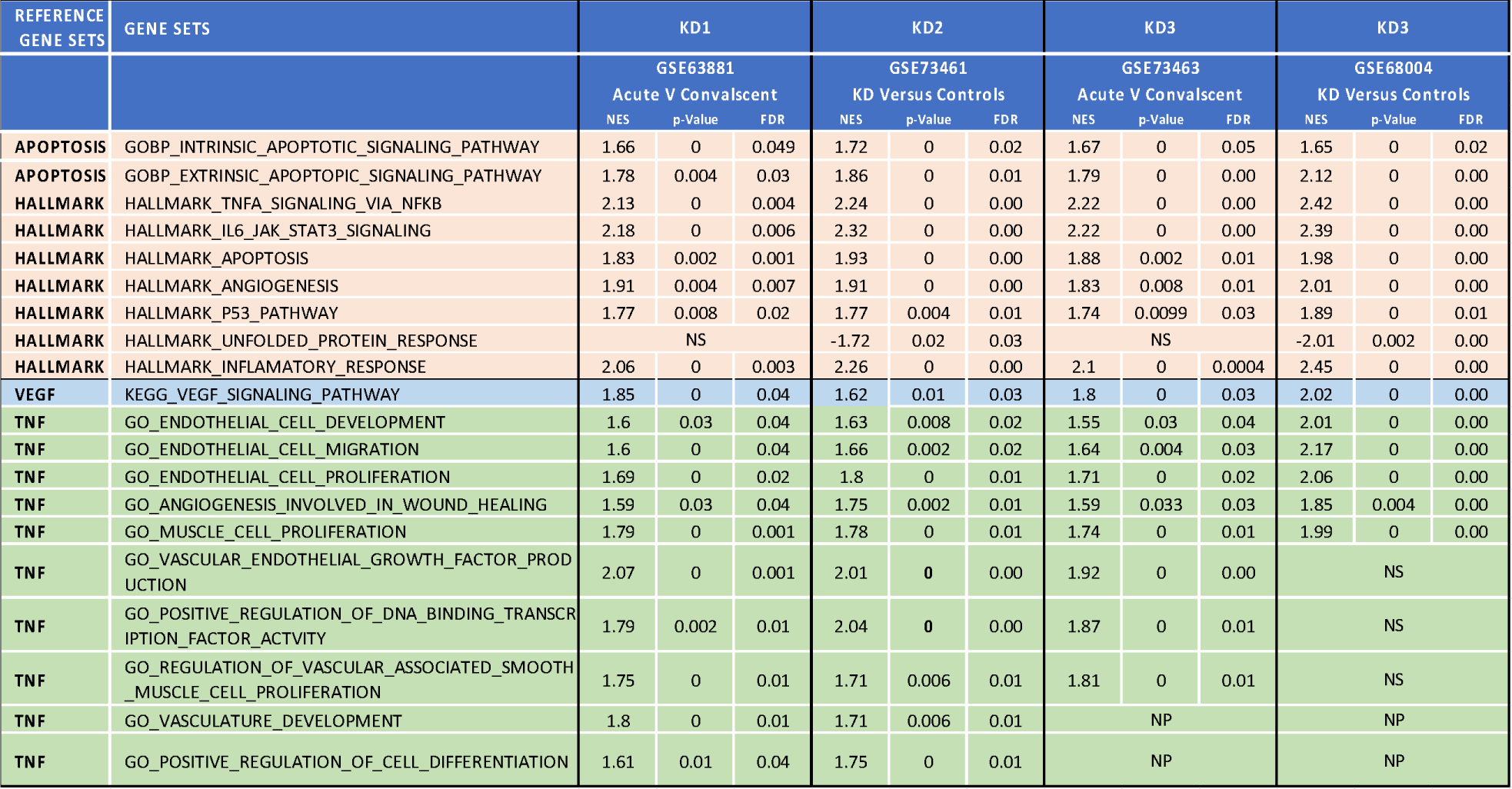
GSEA summary of three datasets. **Table 2**. NES= Net Enrichment score, p-Value and FDR = false discovery rate, NS = Not significant. A two-group t-test was undertaken in each of the KD datasets (KD1-KD3) with a p<0.05 and q(FDR)<0.05. Different reference gene sets were used for pathway analysis as shown in the leftmost column. In KD1 (GSE63881) Acute (n=171) versus convalescent (n=170) samples were compared. For KD 2 (GSE 73461), KD cases (n=78) were compared against 55 (afebrile) controls. Also, for KD3 (GSE68004), KD (n=89) was compared against controls (n=37). If, after analysis the pathway did not appear on the list of genes tested by GSEA, this was marked as Not Present (NP). Gene pathways are shown with the corresponding statistical parameters, including the Normalised Enrichment Score (NES). A negative NES suggests downregulation of the gene set pathway.

Using the Hallmark gene sets for GSEA, the TNFA signaling via NFKB pathway was found to be significant in all three datasets ([KD1:p=0,q=0.003][KD2:p=0,q=0.00] [KD3:p=0,q=0.00]), which supports the presence of an inflammatory response in KD. IL6 JAK STAT 3 signaling was also found to be significant in all three datasets ([KD1:p=0,q=0.006][KD2:p=0,q=0.00] [KD3:p=0,q=0.00]). Additionally, for KD1-KD3, intrinsic (p=0,p=0,p-0) and extrinsic (p=0.004,p=0,p=0) apoptotic signaling were also found to be enriched. The Angiogenesis pathway, as defined by the Hallmark and GO databases, was also enriched in all three datasets, along with VEGF signaling.

However, the Unfolded Protein Response pathway was found to be enriched only in the KD2 and KD3 datasets where KD was compared to controls. The GO pathways enriched for KD1-KD3 datasets in relation to the Endothelium included those related to development, migration, and proliferation, suggesting endothelial activity within vasculature in KD. Enriched GO pathways for vascular development and vascular-associated smooth muscle proliferation further support these findings.

Therefore the actin reference set was also used to compare samples before and after treatment (KD1) and in KD versus controls (KD2-3). GO pathways for various aspects of the actin molecular pathways are shown (Table 4). Enriched pathways included those related to Actin organization and biogenesis, cytoskeleton reorganization, filament-associated severing, and movement. Also, the leucocyte transendothelial migration pathway was noted to be significant across all three datasets.

**Table 4:** GSEA KD1, KD2 and KD3 for Actin associated pathways. **Table 2** Actin pathways are explored in three separate clinical studies (KD1, KD2, and KD3) with KD compared against. GSEA was generated using the Actin GMT (downloaded from the MSigDB) as the reference gene set (left-most column). NES= Net Enrichment score, p-Value and FDR = false discovery rate, NS = Not significant. The table includes significant pathways as generated (p<0.05 and q<0.05). Gene pathways are shown with the corresponding statistical parameters including the Normalised Enrichment Score (NES). For the KD1 dataset, KD cases before and after treatment were compared, and for KD2 and KD3, KD cases were compared against controls.

| REFERENCE GENE SETS | GENE Pathways | KD1 |  |  | KD2 |  |  | KD2 |  |  | KD3 |  |  |
| --- | --- | --- | --- | --- | --- | --- | --- | --- | --- | --- | --- | --- | --- |
|  |  | GSE63881 |  |  | GSE7361 |  |  | GSE7363 |  |  | GSE68004 |  |  |
|  |  | Acute Versus Convalescent |  |  | KD versus Controls |  |  | Acute Versus Convalescent |  |  | KD versus Controls |  |  |
|  |  | NES | p | q | NES | p | q | NES | p | q | NES | p | q |
| Actin | Actin Cytoskeleton Organisation and Biogenesis | 1.66 | 0.02 | 0.03 | 1.73 | 0.006 | 0.01 | 1.63 | 0.014 | 0.04 | 2.13 | 0 | 0 |
| Actin | GOBP Actin Cytoskeleton reorganisation | 1.72 | 0.004 | 0.02 | 1.75 | 0.002 | 0.01 | 1.62 | 0.01 | 0.042 | 1.98 | 0 | 0 |
| Actin | GOBP Actin Filament Based Movement | 1.81 | 0 | 0.01 | 1.77 | 0 | 0 | 1.82 | 0 | 0.01 | 1.95 | 0 | 0 |
| Actin | GOBP Actin Filament Depolymerization | 1.76 | 0.002 | 0.01 | 1.93 | 0 | 0.002 | 1.64 | 0.013 | 0.038 | 1.98 | 0 | 0 |
| Actin | GOBP Actin Filament Organisation | 1.92 | 0 | 0.004 | 1.95 | 0 | 0 | 1.85 | 0 | 0.009 | 2.29 | 0 | 0 |
| Actin | GOBP Actin Filament Severing | 1.91 | 0.002 | 0 | 2.02 | 0 | 0 | 1.82 | 0.006 | 0.01 | 2.06 | 0 | 0 |
| Actin | KEGG Leucocyte Transendothelial Migration | 2.01 | 0 | 0 | 2.08 | 0 | 0 | 2 | 0 | 0.008 | 2.29 | 0 | 0 |

### F. Bulk tissue expression of VEGF-A

Using the GTEx biobank portal, a profile of VEGF-A was generated across tissues and also specifically for the heart (Figure 4). This shows the high propensity of VEGF-A expression in the myocardium compared to other tissues. Further in the heart itself, VEGF-A is expressed highly in the atrial appendage, aorta, and adipose tissue. VEGF-B is also expressed in adipose tissue, though showing lower overall Transcripts Per Million (TPM) compared to VEGF-A.

**Figure 4.**
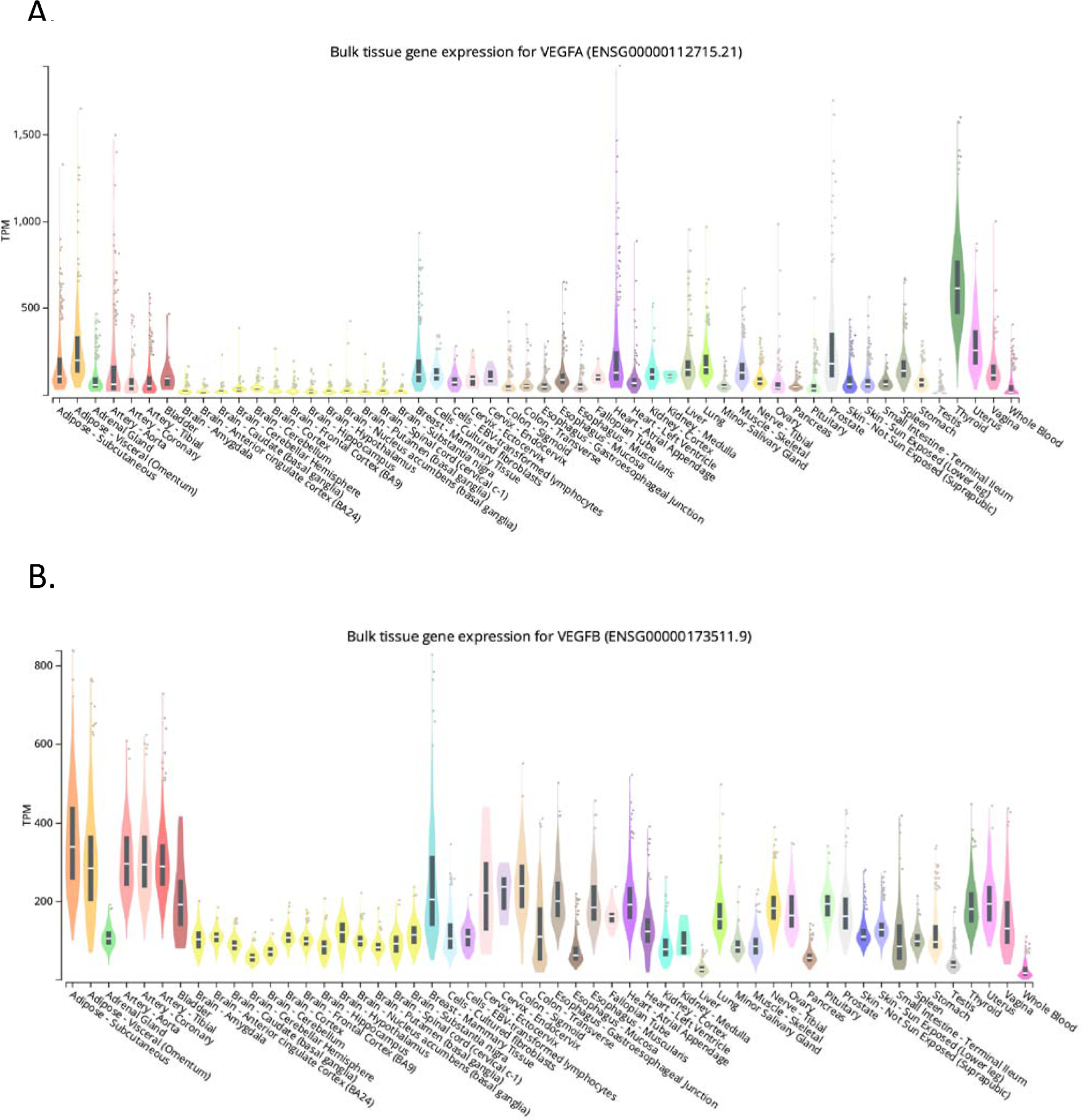
Bulk tissue expression of VEGF-A and VEGF-B a Global Representation. **Figure 4.** Bulk tissue expression studies are shown, generated from the GTEx biobank portal. Expression values are shown in TPM (Transcripts Per Million), calculated from a gene model with isoforms collapsed to a single gene. Box plots are shown as median, 25^th^ and 75^th^ centimes, points are displayed as outliers if they are above or below 1.5 times the interquartile range. Bulk tissue expression of VEGFA in global tissue database showing high median TPM values in cardiac tissues indicating gene expression may impact the severity in KD associated with coronary artery disease (Figure 4A). Bulk tissue expression of VEGFA in global tissue database showing high median TPM values in cardiac tissues indicating gene expression may impact the severity in KD associated with coronary artery disease. Gene expression for VEGF-B is also shown (Figure 4B). The data is not age-dependent so the comparison between children and adults is not possible.

### G. Gene Function network representation of VEGF and HSP genes

Through the GeneMania platform, a functional network association representation of VEGF and HSP genes is presented (Figure 5). The genes HSPBAP1, VEGF-A and VEGF-B are shown to be highly connected.

**Figure 5.**
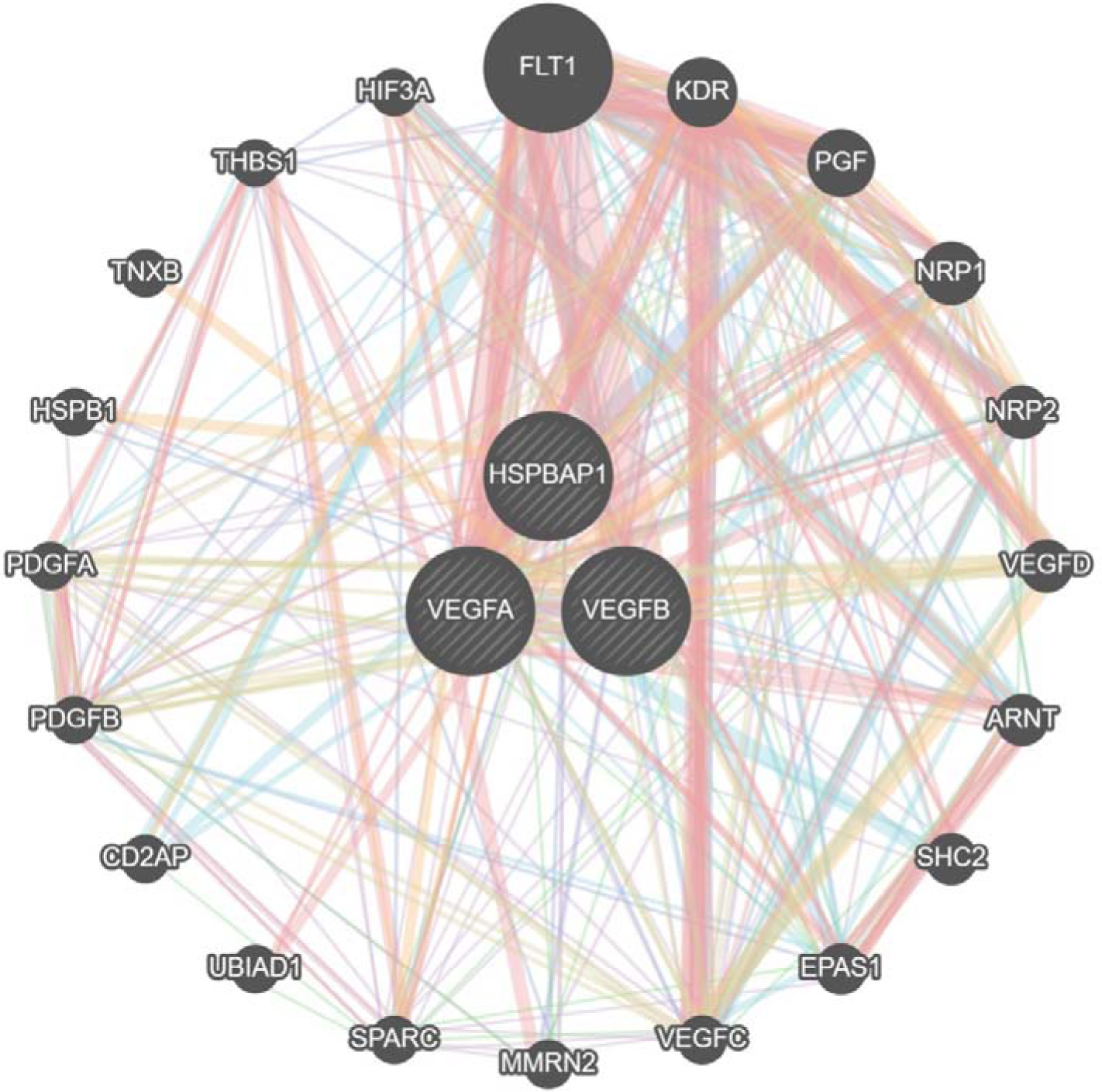
Gene functional Network Association of VEGF and Heat Shock Proteins. A functional gene network association representation is shown generated from the GeneMania portal by inserting the genes shown in the illustration. Gene Function Interaction among VEGF-A, VEGAF-B and HSPBAP1 was observed to have a greater physical and co-expression interaction signifying that gene interaction may play an important role in KD pathogenesis.

**Figure 6.**
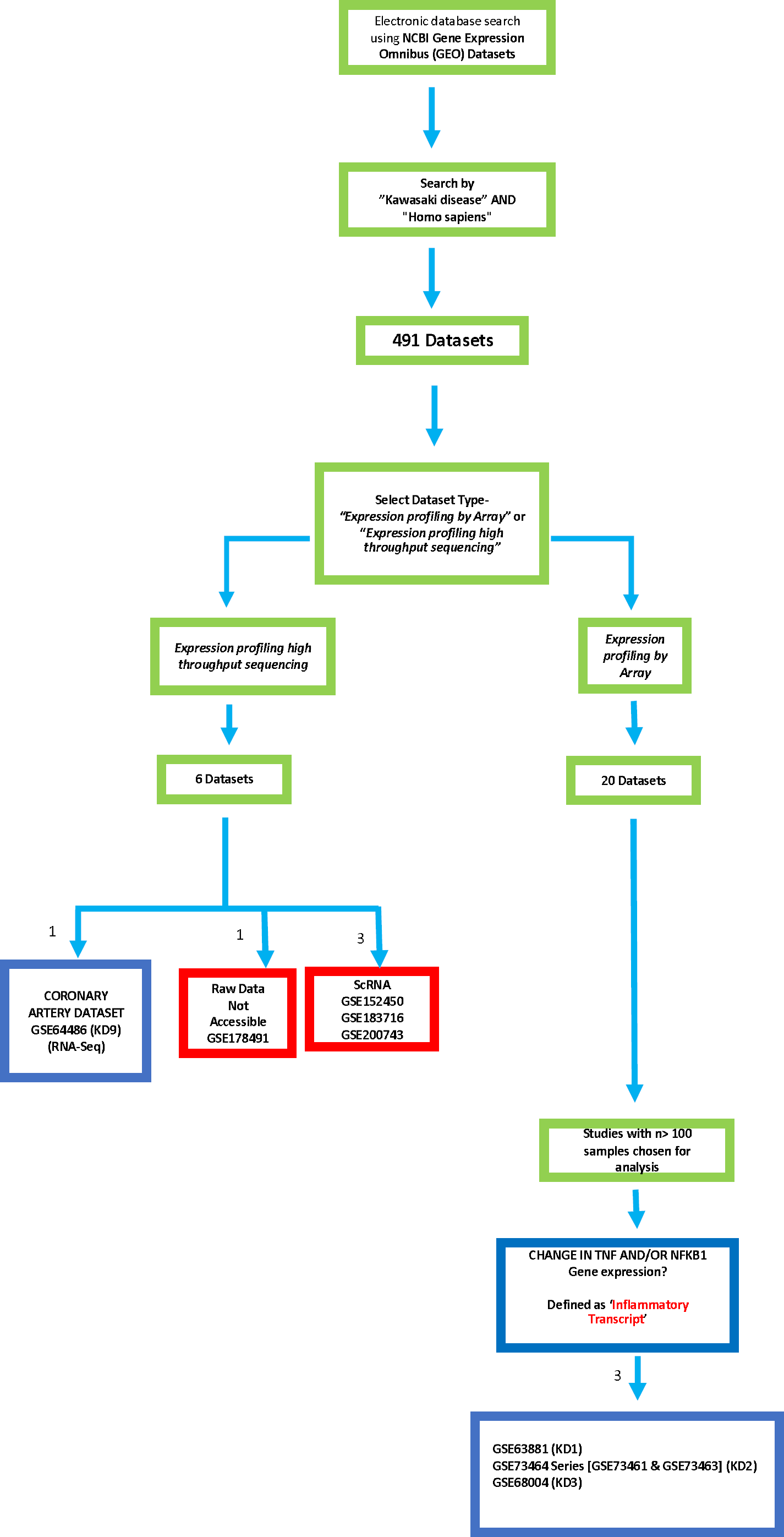
Systematic Search for KD datasets from DNA Microarray and RNASeq datasets. Discriminate Molecular Mechanisms between KD positive and the Controls. **Figure 6.** A systematic search for KD datasets was undertaken from the NCBI GEO database. The different stages in the filtering and selection process are shown, with the number of datasets at each filtering stage indicated next to the direction arrows. The search term “Kawasaki Disease” and “Homo Sapiens” generated 491 datasets. This was then divided into “Expression Profiling by Array (20 datasets) and Expression profiling by high throughput sequencing (6 Datasets). Of the 6 datasets, the datasets shown in red were excluded due to not being accessible, inadequate labeling, or entailing sc-RNA studies, or were non-KD studies; which left one dataset (GSE64486) for analysis. GSE64486 is a unique dataset involving coronary artery tissue as we wanted to include the larger studies, a n of > 100 was chosen, which lead to the filtering of the 20 datasets down to three datasets (GSE63881, GSE73464 and GSE68004). GSE73464 is the super series containing sub-series GSE73461, GSE73462, and GSE73463.

## Discussion

Systemic inflammation in KD places children at risk of coronary arteritis and aneurysm formation during the acute febrile period. Our analysis aimed to understand the underlying inflammation-associated processes, assuming the presence of a thermal stress response related to HSPs and VEGF. Transcriptomic analysis was conducted using three peripheral blood datasets (KD1-KD3) and one coronary artery dataset (KD4). KD1 and KD2 compared acute versus convalescent samples, while KD2, like KD3, also included KD versus control samples. To track inflammation, we examined the gene expressions of tumor necrosis factor (TNF) and nuclear factor kappa B1 (NFKB1). Analysis of peripheral blood samples from KD1-KD3 revealed a significant increase in TNF and NFKB1 gene expression, suggesting inflammation based on gene expression profiles. Gene set enrichment analysis (GSEA) of KD1-KD3 datasets identified activation of the Hallmark IL6 JAK STAT3 signaling pathway^13^. This pathway is involved in inflammation, immune response, and cell growth in across various conditions. Our study of KD1-KD3 revealed GSEA enrichment of gene sets associated with endothelial cell development, migration, proliferation, leukocyte transendothelial migration, vascular changes, angiogenesis, intrinsic and extrinsic apoptotic signaling, pathways related to vascular development, providing insights into the involvement of endothelial and angiogenic effects in KD-induced inflammation. The coronary artery dataset (KD4), on the other hand, showed no statistically significant difference in gene expression, potentially indicating end-stage disease with diminished inflammation.

HSPs play a crucial role as chaperones, assisting in the correction of misfolded proteins. One question of interest is how HSPs are triggered during KD-associated inflammation. To investigate this, we examined the unfolded protein response (UPR) pathway in KD. The UPR is a cellular stress response activated by the accumulation of misfolded or unfolded proteins ^14^. Gene set enrichment analysis (GSEA) applied to the transcriptomic data revealed the enrichment of hallmark gene ontology (GO) pathways, including the UPR pathway, in two datasets. Activation of HSPs, such as HSP70, can induce the UPR, leading to either apoptosis to eliminate pathogenic cells or survival by preventing apoptosis in host cells. Furthermore, the UPR plays a significant role in angiogenesis by modulating VEGF-A. Recent studies have shown that p53, a heat shock protein 90 (HSP90), is involved in the UPR ^15^ and is crucial in coordinating immune responses to infection. GSEA also showed the p53 pathway was enriched in KD patients compared to controls (KD1-KD3). Notably, p53, as an HSP, may play a significant role in the inflammation observed in various conditions such as KD and sepsis. Further, Wong et al. (2017) developed a biomarker protein panel for sepsis endotyping, which included p53 in the prediction matrix ^16^. HSPBAP1, a relatively uncharacterized gene, is suggested to contribute to the control of the HSP27 pathway, which is important for cellular responses to various stressors, including heat shock, oxidative stress, and inflammation. This study found HSPBAP1 gene expression significantly distinguished KD from control samples. The predicted gene function analysis of VEGF and HSPBAP1 revealed substantial physical interaction (>70%), suggesting genetic mechanisms may influence risk attribution and interactions with other pathway genes. Further investigation is needed to elucidate HSPBAP1 function and its interactions within the KEGG pathway.

We observed the consistent up-regulation of pro-inflammatory genes VEGF-A, TNF, and NFKB1 in the Acute phase of KD and when KD samples were compared to controls (KD1-KD3). Given their temporal relationship, we termed these genes the Temporal Transcript Model (TTM). In this paper, our analysis focused on specific HSPs reported to be associated with angiogenesis, namely HSPB1, HSPA1A, and HSP90AB1 (Table 1). A similar pattern of GE up-regulation was noted for HSPA1A, whereas HSP90AB1 GE was observed to be down-regulated in the TTM (KD1-KD3). Molecular chaperones HSP70 and HSP90 play a role in protein remodeling. HSPB1, a small heat shock protein, is involved in both angiogenesis and apoptosis^17^. It interacts with VEGF-A, a crucial regulator of angiogenesis ^18^, stabilizing and increasing the number of VEGF receptors on the cell surface, thereby enhancing VEGF signaling and promoting angiogenesis ^19^. Additionally, the interaction between HSPB1 and p53 may play a role in cellular protection against stress and damage. HSPB1 interacts with p53, increasing HSPB1 stability and enhancing its regulation of cell cycle and apoptosis. HSPs, such as HSP70 and HSP90, play a regulatory role in the activity and expression of various growth factors, including vascular endothelial growth factor (VEGF) ^20^. Through binding to VEGF receptors and influencing its signaling pathways, HSPs modulate VEGF activity, thereby affecting angiogenesis and blood vessel formation. The relationship between NF-kappa B activated inflammation, HSPs, and VEGF is exemplified in a human microvascular tissue-engineered endothelial model ^21^, which demonstrated activation of the Heat Shock Protein (HSP) and NF-kappa B pathways along with VEGF production and capillary formation. Moreover, in a human model, an elevation in body temperature was associated with a transient increase in mRNA gene expression of both VEGF and HSP, indicating a potential link between these two protein families ^22^. Based on this, we hypothesized that changes in gene expression of angiogenesis-inducing HSPs, VEGF-A, and VEGF-B, are associated with coronary artery disease (CAD) in KD.

VEGF-A gene expression was found to be increased in acute versus convalescent samples and in Kawasaki disease (KD) patients versus controls, suggesting a potential role in KD progression. In contrast, VEGF-B gene expression decreased in the same comparisons, indicating a potential counter-regulatory mechanism. Gene set enrichment analysis (GSEA) revealed the enrichment of gene ontology (GO) pathways related to VEGF production, supporting the involvement of VEGF signaling in KD pathogenesis. Analysis of the global tissue expression database, which provides bulk tissue gene expression data (gtexportal.org), showed high VEGF-A transcript per million (TPM) values in the heart and high VEGF-B TPM values in arterial tissue. This observation implies that changes in VEGF-A or VEGF-B expression levels may significantly impact the cardiovascular system in KD patients. To better understand the specific roles of VEGF-A and VEGF-B in KD and their potential influence on disease progression and cardiovascular complications, further investigation is required. This research should include the examination of age-related changes in VEGF-A and VEGF-B expression and their subsequent cardiological effects in the context of KD. HSPB1 may have a role as a regulator of actin filament dynamics^23^. The complex process of coronary aneurysm formation in KD involves various factors, including changes in the mechanical properties of the vessel wall, blood flow patterns, and growth factors and cytokines that promote inflammation, leading to weakness in the vessel wall ^24^. Structural changes affecting the coronary vessels are key contributors to aneurysm formation, and the TTM GSEA demonstrated alterations in actin-based pathways. These effects play a crucial role in the development of coronary artery lesions in KD.

Additionally, matrix metalloproteinases (MMPs), zinc-dependent extracellular matrix endopeptidases involved in remodeling, have been shown to contribute to VEGF-A generation ^17, 25^. MMPs are involved in various physiological processes, including morphogenesis, embryogenesis, angiogenesis, and wound repair. In the context of KD, MMPs may play a role in the structural changes of coronary arteries during aneurysm formation. In a murine model of KD, elevated levels of NFKB and MMP9 were observed in the coronary artery trunk and branches, with no apparent broken elastin observed in the IVIG and control group, and a less severe inflammatory cell infiltrate noted ^26^. These findings suggest that MMPs and NFKB may contribute to the structural changes of coronary arteries during aneurysm formation in KD. Targeting the upregulation of these genes in the coronary arteries could be a potential therapeutic approach to reduce inflammation and prevent aneurysm formation. Furthermore, recent research has emphasized the significance of endothelial cell response in KD pathogenesis. Endothelial cells play a critical role in maintaining vascular integrity and regulating the balance between pro-and anti-inflammatory factors. In KD, activation and injury of endothelial cells are believed to contribute to the development of coronary artery aneurysms.

This paper has two main limitations: firstly, the arbitrary study selection parameter, and secondly, the choice of genes examined from a differential gene expression (GE) perspective. An arbitrary threshold of 100 samples was employed for selecting key studies for secondary analysis, prioritizing statistical power, reproducibility, effect size estimation, generalizability, reduction in sampling bias, and reliable identification of differentially expressed genes. This strategy resulted in the identification of statistically repeatable GE patterns. However, the choice of KD1-KD3 is consistent with our earlier work which selected the same datasets as showing a consistent temporal change in gene expression with respect to the genes under study in KD. The relationship between VEGF-A, HSPs, and coronary endothelial cells (end-point) has not been extensively studied in the context of KD. Therefore, when constructing a suitable temporal transcript framework for KD, genes of interest were assumed based on their pro-inflammatory function. VEGF-A was included due to its central role in angiogenesis, and VEGF-B was considered due to its competition for VEGFR-1 receptor binding with VEGF-A. This framework enabled the examination of changes in VEGF and HSP gene expression using a systems approach offered by gene expression analysis. However, additional research is necessary to substantiate the downstream effects on molecular pathways and to investigate the functional implications of these gene expression changes. Despite these limitations, our comprehensive approach offers researchers an overview of key pathways and valuable insights across large datasets.

Studies suggest that actin dynamics play a crucial role in aneurysm development and progression. GSEA was used to compare pre-and post-treatment samples (KD1) and KD versus control samples (KD2-3) using the actin reference set, revealing enriched pathways related to actin organization, biogenesis, cytoskeleton reorganization, filament-associated severing, and movement. Additionally, the leukocyte transendothelial migration pathway was significant. Actin dynamics involve the regulated assembly and disassembly of actin filaments, while aneurysm formation is characterized by arterial wall weakening and bulging. Dysregulated actin dynamics in smooth muscle and endothelial cells can contribute to aneurysm-forming arterial wall changes. Suzuki et al. (2000) observed extensive vascular growth factor expression in thickened intima at stenotic sites and recanalized vessels in KD patients^27^. Active coronary artery lesion remodeling in KD, including intimal proliferation and neoangiogenesis, continues for years after disease onset. The ongoing process beyond the acute phase could explain why GSEA did not distinguish acute from convalescent samples based on UPR, while there was a difference between KD and control samples. Further research is needed to explore ongoing molecular changes in KD.

The temporal studies conducted in this paper demonstrate the value of considering gene expression profiling across time and the usefulness of end-stage transcriptional datasets as a cross-sectional tool. Understanding the modulation of the pyrexial response in KD to prevent coronary artery lesions could be particularly valuable. Future studies should also take into account the HSP response when assessing treatment efficacy, especially in modulating strategically important molecules like HSPB1. The TTM framework we constructed offers a valuable foundation for future KD research, particularly with regard to age and ethnicity considerations. The transcriptomic approach in this paper offers a comprehensive view of multiple processes, providing an integrated understanding of complex biological mechanisms.

## Conclusion

The transcriptional analysis of key KD datasets sheds light on the complex interplay between VEGF-HSP-associated gene expression, inflammation, and angiogenesis in KD pathogenesis. The study identified changes in the gene expression of specific HSPs associated with angiogenesis and demonstrated that HSPBAP1 gene expression significantly distinguished KD from control samples. HSPs, especially HSPB1, play a role in thermal injury and inflammation, while altered VEGF-A and VEGF-B gene expression is associated with KD-associated angiogenesis. Gene set enrichment analysis (GSEA) revealed enrichment of gene ontology (GO) pathways related to VEGF production. The relationship between NF-kappa B-activated inflammation, HSPs, and VEGF is exemplified in a human microvascular tissue-engineered endothelial model. Further research is needed to elucidate the precise role of HSPs in KD and explore its potential as a therapeutic target, as well as to investigate the interplay between HSPs associated with angiogenesis, VEGF-A, and VEGF-B in KD. The transcriptomic approach offers a systems-level perspective, providing an integrated understanding of complex biological processes. The temporal transcript framework constructed offers a valuable foundation for future KD research. It suggests a framework based upon key KD datasets in the scientific literature useful for hypothesis testing and planning suitable studies in the future.

## Notes

https://www.ncbi.nlm.nih.gov

### Competing Interest Statement

The authors have declared no competing interest.

### Summary of Updates

Figures have been updated, with editing of text

